# Enteric infection priming confers IL-17A–dependent protection from chemically-induced Colitis

**DOI:** 10.1101/2025.05.03.652030

**Authors:** Vishwas Mishra, Rita Berkachy, Priyanka Biswas, Gad Frankel

## Abstract

**Background and Aims:** Enteric infections trigger mucosal immune responses. However, whether such immune imprinting influences susceptibility to sterile inflammatory diseases like colitis remains unclear. The aims of this study were to investigate whether a resolved *Citrobacter rodentium* (CR) infection in mouse alters host susceptibility to chemically induced colitis and to identify the underlying immune mechanisms.

**Methods:** C57BL/6 mice were infected with wild-type CR or CR Δ*map*Δ*espF*, a mutant lacking tight junction-disrupting effectors. 3 weeks post clearance, mice were subjected to dextran sodium sulfate (DSS)-induced colitis. Disease severity was assessed by weight change, colon length, histopathology, and myeloperoxidase levels. Colonic immune cell populations were characterised by flow cytometry, and cytokine levels were measured from colon explants. Functional roles of IL-17A were evaluated using recombinant cytokine administration and neutralising antibody treatment.

**Results:** Mice that cleared CR infection fared better following DSS treatment compared to uninfected or Δ*map*Δ*espF*-infected mice. This protective phenotype was not directly dependent on microbiota, as confirmed by co-housing experiments. Protected mice displayed elevated numbers of colonic Th1 and Th17 cells and higher levels of IL-17A, IL-22, and IL-2 cytokines. Prophylactic treatment with IL-17A conferred protection in naïve mice, whereas IL-17A neutralisation in previously infected mice abrogated the benefit, identifying IL-17A as a key mediator of protection.

**Conclusions:** Resolved intestinal infection with CR confers long-term protection against colitis via persistent IL-17A-mediated immune reprogramming. These findings resonate with the “hygiene hypothesis” and highlight how prior microbial exposure can shape mucosal resilience.

## Introduction

Intestinal mucosal injury can be caused by diverse triggers, including infection and chronic inflammation, such as inflammatory bowel disease (IBD)^1^. While the initiating causes differ, both infectious and non-infectious colitis converge on common pathological features, including epithelial barrier disruption, cytokine-driven inflammation and immune cell infiltration^1,2^. This convergence raises the possibility that immune responses shaped during enteric infections may influence how the host responds to ensuing mucosal perturbations. However, whether infection-induced immune imprinting modulates susceptibility to subsequent inflammatory disorders such as IBD, remains poorly understood.

*Citrobacter rodentium* (CR) is an extracellular enteric murine pathogen, which shares infection strategies with the human attaching and effacing (A/E) pathogens enteropathogenic and enterohemorrhagic *Escherichia coli*, has served as a powerful tool for dissecting host– pathogen interactions at the mucosal interface^2,3^. Adhesion of A/E pathogens to intestinal epithelial cells (IECs) is mediated by injection of bacterial effector proteins by a type III secretion system, including Tir, which mediates intimate bacterial attachment to the apical surface of IECs and EspF and Map, which disrupt tight junctions (TJs)^4^. Infection with CR induces colonic crypt hyperplasia (CCH), disruption of the intestinal barrier, inflammation, diarrhoea and reprogramming of host metabolism^2,3^. CR infection leads to significant changes in the gut microbiome, characterised by reduced microbial richness and distinct alterations in specific bacterial taxa^5,6^. CR infection in C57BL/6 mice can be divided into four phases: Establishment (1–3 days post infection (dpi)), where CR colonises the caecal lymphoid patch. Expansion (4–8 dpi), characterised by rapid CR proliferation and adherence to IECs in the distal colon with disruption of TJ; Steady-State (8–12 dpi), characterised by high CR shedding and stable colonisation; Clearance (12-18 dpi), CR is rapidly cleared from 13 dpi^3^.

During the expansion phase CR triggers innate immune responses, characterised by secretion of IL-17A and IL-22 from group 3 innate lymphoid cells (ILC3), IL-18 from IECs and recruitment of neutrophils^2,7^. The clearance phase is characterised by expansion of IgG-producing B cells and CD4⁺ T cells, with key roles played by Th1 cell producing TNF, IL-2 and IFNγ and Th17 cells secreting IL-17A/F, and IL-22—cytokines essential for maintaining barrier function, controlling bacterial growth, and limiting systemic dissemination^2,7^. These immune responses not only contribute to pathogen clearance but also leave a lasting imprint on the mucosal immune landscape. Previous studies have shown that ILC3s and resident memory T cells persist in the colon months after resolution of CR infection, which protect against reinfections^8,9^. Moreover, IFNγ signatures also persist weeks post CR clearance^10^. However, whether these infection-primed immune responses can affect outcomes against future unrelated, sterile inflammatory insults remains largely unexplored.

IBD, encompassing Crohn’s disease and ulcerative colitis, are chronic, inflammatory, relapsing disorders of the gastrointestinal tract^1^. Although the precise causes remain elusive, IBD is widely understood to arise from a multifactorial interplay between host genetics, environmental factors, altered microbiota, and immune dysregulation^1^. Central to IBD pathogenesis is an imbalance in mucosal immune responses, including exaggerated activation of effector T cells and disruption of cytokine networks that normally maintain epithelial barrier integrity^11^. Among the type 3 cytokines implicated, IL-22 plays a key protective role in intestinal homeostasis by acting directly on IECs to promote wound healing, strengthen barrier function, and induce antimicrobial peptide expression^12^. Its beneficial effects are well documented in murine models of colitis, where IL-22 deficiency exacerbates disease severity, while recombinant IL-22 treatment ameliorates inflammation^13^. In contrast, the role of IL-17A in IBD remains controversial^14,15^. While IL-17A contributes to barrier maintenance and mucosal defence in some contexts, clinical trials targeting IL-17A have worsened colitis symptoms in patients, revealing a tissue- and disease-specific complexity in its function^14^. These observations underscore the need to better define the conditions under which IL-17A acts as a friend or foe in intestinal inflammation.

Dextran sodium sulfate (DSS)-induced colitis is one of the most widely used experimental systems to model aspects of ulcerative colitis^16^. DSS administration in drinking water causes direct chemical injury to colonic epithelial cells, leading to barrier disruption, microbial translocation, and activation of innate and adaptive immune responses, recapitulating several features of acute intestinal inflammation observed in patients^17^. The DSS model has been used to study the role of epithelial repair mechanisms, cytokine responses, and mucosal resilience following barrier disruption^16,17^.

In this study, we investigated whether a history of resolved CR infection alters susceptibility to DSS-induced colitis. Specifically, we focused on the long-term changes in the colonic immune landscape—particularly T cell responses and cytokine production. This uncovered a novel role for infection-induced IL-17A in protecting the colon from subsequent perturbation following CR clearance. Our work highlights the importance of immune response training by enteric infection for protecting the gut mucosa from chronic inflammatory disorders.

## Material and methods

### Mouse experiments

All animal work was conducted at Imperial College London (Association for Assessment and Accreditation of Laboratory Animal Care accredited unit) under the auspices of the Animals (Scientific Procedures) Act (UK) 1986 (PP7392693). The animal work was approved locally by the institutional ethics committee. Pathogen-free 18–20-g female C57BL/6 were purchased from Charles River Laboratories and housed in groups of 5 in dedicated animal facilities of Imperial College London (12h light/dark cycle; 22 ± 2°C; 30 to 40% humidity). Mice were housed in IVC cages with corn cob bedding and enrichments including refuges, nesting material, and gnawing sticks. Mice were fed with RM1(E) rodent diet (SDS Diet) and water ad libitum.

### CR infection and DSS-induced colitis

CR was grown overnight in lysogeny broth containing 50 μg/mL nalidixic acid at 37°C with shaking at 180 rpm, centrifuged at 3000 x g for 10 min and resuspended in sterile PBS^18^. Mice were infected with approximately 1 x 10^9^ CFU in 200 μL sterile PBS using oral gavage. Mock infected (PBS) mice received 200 μL sterile PBS. The inoculum CFU was retrospectively confirmed by CFU quantification. Mice were monitored every day. Body weight and CFU counts were assessed as previously described^18^. Briefly, analysis of CFUs was determined via serial dilutions of homogenized faecal pellets on the indicated days post CR infection, followed by plating on LB agar plates supplemented with 50 μg/mL nalidixic acid.

At 40 dpi, mice were weighed and then administered with 2.5% DSS (MP Biomedical, catalogue number 9011-18-1) in drinking water for 7 days, followed by clean drinking water for 3 days. Mice were monitored every day for changes in weight and disease severity. For Co-housing experiments, CR infected mice post clearance (∼20 dpi) and mock infected mice (PBS) were co-housed in groups of 5 mice per cage. To confirm no transmission or presence of CR infection in the Co-housed cage, fresh faecal samples were collected on 8 days post Co-housing, homogenised in sterile PBS, serially diluted and plated on LB-agar containing 50 μg/mL of nalidixic acid. 20 days post co-housing, mice were administered with 2.5% DSS in drinking water for 7 days, followed by clean drinking water for 3 days.

To determine the effect of rIL-17A treatment on outcomes of DSS-induced colitis, naïve mice were treated with 2 µg of recombinant IL-17A (BioLegend, catalogue number 576006) intraperitoneally on 6, 4 and 0 days before the start of DSS administration. For anti-IL-17A treatment, CR infected mice post clearance (CR-Ψ) were injected with 100 µg of anti-IL-17A antibody (ThermoFisher Scientific, catalogue number 16-7173-85) intraperitoneally on 6, 4 and 0 days before the start of DSS administration.

### Post-mortem pathophysiological analysis

Mice were humanely euthanised. The large intestine of mouse consisting of caecum and colon was harvested and placed on a clean surface in parallel to a mm scale and the colon length was recorded and captured using a digital camera. Colon was removed from caecum, cleaned and from the distal side of colon, 0.5 cm colon was stored in 4% paraformaldehyde for histological studies, next 0.5 cm was collected for explant culture, next 0.5 cm was snap frozen in dry ice for MPO analysis and 3 cm was collected for FACS analysis.

### Cytokine profiling from explants

A 0.5-cm segment of distal colon, cleared of faecal matter, was weighed and incubated in RPMI 1640 medium supplemented with glutamine (Sigma), streptomycin (100 μg/mL), and penicillin (100 μg/mL) for 2 hours at RT. After this initial incubation, tissue explants were transferred to complete RPMI medium at a concentration of 1 mL per 0.1 g of tissue. The complete medium consisted of RPMI 1640 with glutamine, supplemented with 10% heat-inactivated foetal bovine serum (FBS), 1 mM sodium pyruvate, 100 μg/mL penicillin, 100 μg/mL streptomycin, and 10 mM HEPES. Explants were cultured under standard conditions (37°C, 5% CO₂) for 24 hours. After incubation, the supernatants were collected, centrifuged at 3,000 x g for 10 minutes to remove residual cells and debris, and stored at −80 °C until analysis.

Cytokine concentrations in the explant supernatants were measured using the LEGENDplex assay kit (BioLegend, catalogue number 741044) following the manufacturer’s protocol. Data acquisition was carried out on a FACSCalibur flow cytometer (BD Biosciences), and cytokine levels were quantified using the LEGENDplex data analysis software (BioLegend).

### Histological analysis

Paraformaldehyde-fixed 0.5-cm distal colon samples were processed, paraffin embedded, and sectioned at 5 μm^18^. The sections were then stained with haematoxylin and eosin (H&E). All images were acquired using a Zeiss AxioVision Z1 microscope with a 20X lens objective using an AxioCam MRm camera and processed using Zen 2.3 (Blue version; Carl Zeiss MicroImaging GmbH, Germany).

### MPO ELISA

0.5 cm frozen colon samples were mechanically homogenized in 500 µL of 1X RIPA buffer containing protease inhibitors (Cell Signalling Technology, catalogue number 9806), followed by centrifugation at 3000 x g for 10 min at 4°C. The supernatant was collected, and total protein concentration was estimated using BCA kit (ThermoFisher Scientific, catalogue number 23227). The MPO concentration was determined using a mouse MPO enzyme-linked immunosorbent assay (ELISA) (Bio-Techne, catalogue number DY3667) according to the manufacturer’s instructions. Readings were obtained using a FLUOstar Omega microplate reader (BMG biotech). The amount of MPO estimated was normalised to the protein concentration.

### Isolation of lamina propria immune cells from mouse colon

Lamina propria cells were isolated from mouse colons using a previously established protocol^19^. Briefly, 3-cm segments of distal colon were excised, washed thoroughly, and opened longitudinally. The tissues were then incubated at 37 °C for 20 minutes in a shaking incubator in calcium- and magnesium-free 1X HBSS containing 2% FBS, 10 mM EDTA, and 1 mM DTT to remove the epithelial layer. Following incubation, the cell suspension was centrifuged to separate IECs from the remaining lamina propria tissue. Supernatants containing IECs were discarded, and the residual tissue was subjected to enzymatic digestion in RPMI 1640 medium containing 62.5 μg/mL Liberase, 50 μg/mL DNase I (Sigma-Aldrich), and 2% FBS at 37 °C for 40–50 minutes. The resulting cell suspension was passed through a 100 μm cell strainer to obtain a single-cell suspension for downstream flow cytometry analysis.

### Flow cytometry

For extracellular staining, single-cell suspensions were distributed into 96-well V-bottom plates and incubated for 10 minutes with LIVE/DEAD Fixable Blue dye (diluted in DPBS) to identify and exclude non-viable cells from analysis. Following this, cells were incubated for 20 minutes with Fcγ receptor blocking reagent (BD Biosciences) to prevent non-specific antibody binding, and then stained for surface antigens using fluorochrome-conjugated monoclonal antibodies. All staining procedures were carried out at 4 °C in the dark unless otherwise specified. Unstained controls, LIVE/DEAD-only controls, and fluorescent-minus-one (FMO) controls were included to assess background fluorescence and set gating thresholds. After staining, cells were washed and fixed for 20 minutes using the eBioscience Foxp3/transcription factor fixation buffer set. Fixed samples were stored in the dark at 4 °C until acquisition.

For compensation, single-color controls were prepared using UltraComp eBeads™ Compensation Beads. Prior to analysis, cells were washed and resuspended, and data were acquired on 50,000 live events per sample using an Aurora flow cytometer (Cytek Biosciences). Flow cytometric data were analysed with FlowJo software (version 10.8.1, Tree Star). The gating strategy used to identify various T cell subsets is illustrated in Figure 4A.

### Statistical analysis and data plotting

Sample size was not predetermined using statistical methods. Experimental design followed a randomised block structure, typically including 5 animals per group per experiment, with each experiment independently replicated at least 2 times. Data from all animals across replicates were combined for analysis. Results are presented as mean ± SEM.

For comparisons between two groups, statistical significance was assessed using the student’s t-test. When comparing more than two groups, analysis of variance (ANOVA) was applied. To account for multiple comparisons, a false discovery rate (FDR) threshold of 5% (Q = 0.05) was used, employing Bonferroni correction as implemented in GraphPad Prism version 10.4.0. P values adjusted using FDR that exceeded 0.05 were considered not significant (ns). All data visualisation and statistical analyses were carried out using GraphPad Prism (version 10.4.0, GraphPad Software Inc.). Specific statistical parameters for each experiment are provided in the corresponding figure legends.

## Results

### A history of CR infection protects mice from subsequent chemical-induced colitis

To determine if a history of intestinal infection influences the susceptibility to colitis, we orally infected C57BL/6 mice with ∼ 10^9^ CFU of CR, control mice received sterile PBS. Temporal faecal quantification revealed that CR was cleared by 21 dpi (Fig. S1A-B). At 40 dpi (i.e. 19 days post clearance) the experimental and control mice were treated with 2.5% DSS in drinking water for 7 days, followed by 3 days of clean drinking water (7+3 model) (Fig. 1A). DSS treatment resulted in disease as evidenced by weight loss (Fig. 1B). Mice with a history of CR infection (CR-Ψ, the symbol ‘Ψ’ represents memory/history of infection, hereafter) showed significantly higher weight gain post DSS treatment than mice with no history of infection (PBS) (Fig. 1B).

**Figure 1.**
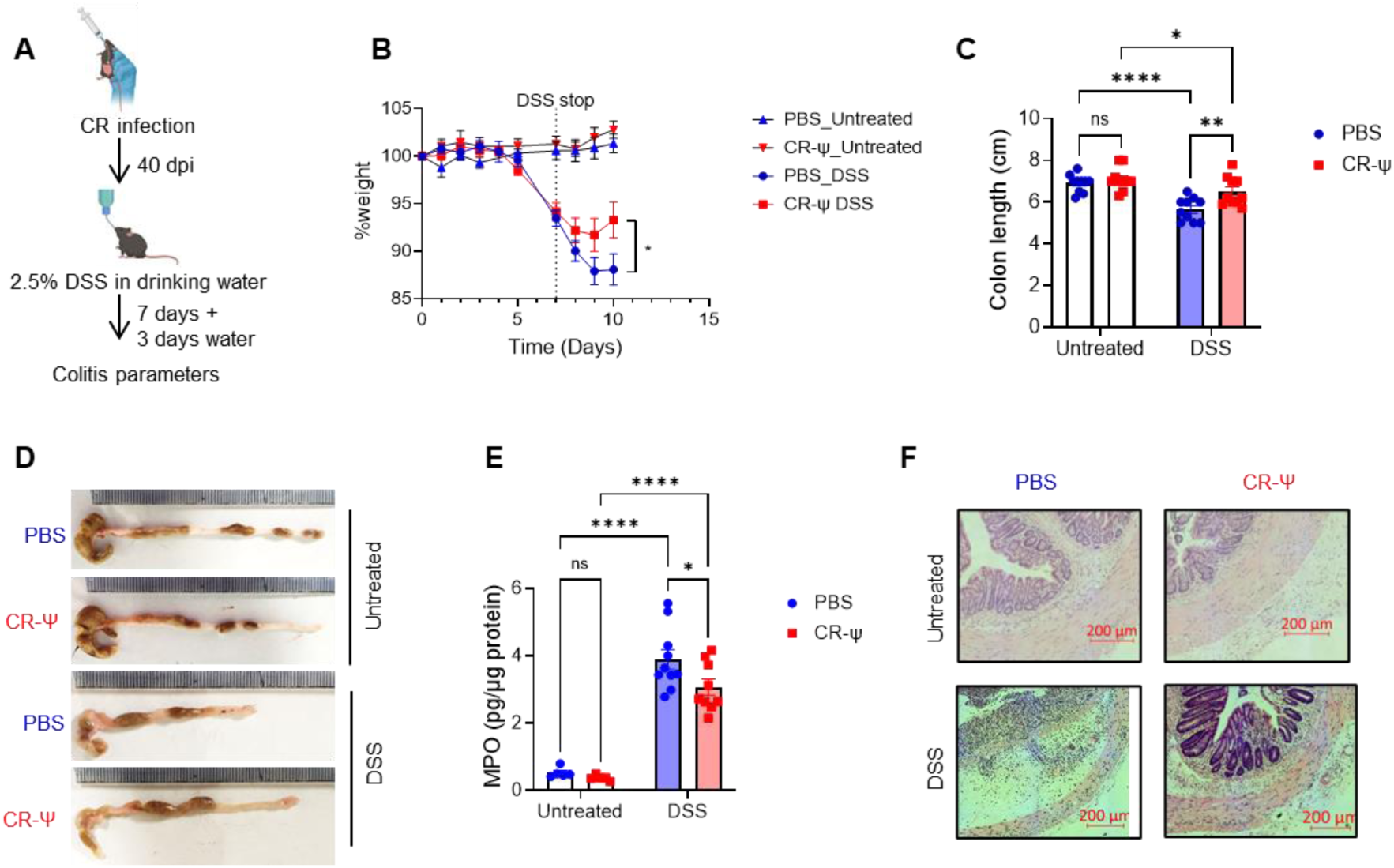
A history of CR infection protects mice from DSS-induced colitis. (A) Schematic representation of the experimental design: C57BL/6 mice were infected with CR or mock-treated with PBS, followed by 2.5% DSS in drinking water at 40 dpi for 7 days, with 3 days of clean water. (B) Weight loss, (C) Colon length measurements post-DSS treatment, (D) Representative colon images, (E) Colonic myeloperoxidase (MPO) levels, a marker of neutrophil infiltration. (F) Representative H&E-stained colon sections show reduced epithelial disruption and immune cell infiltration in CR-Ψ mice. Scale 200 µm. Data shown are pooled values from 2 biological repeats with 5 mice in each group per experiment. P values were determined on data plotted as mean ± SEM using Student’s-t-test in (B) and Two-way ANOVA with Bonferroni post-test for multiple comparisons (C, E). *p < 0.05; **p < 0.01; ****p < 0.0001.

Colonic shortening and faecal myeloperoxidase (MPO) are markers for colitis severity in mice^16,20^. Similar colon length and faecal MPO were recorded in untreated PBS and CR-Ψ mice (Fig. 1C-E). However, while significant shortening of colon was observed in both treatment mouse groups post DSS treatment, significantly longer colon was seen in CR-Ψ compared to PBS mice (Fig. 1C-D). Moreover, the levels of colonic MPO were markedly reduced in CR-Ψ compared to control mice (Fig. 1E).

Histological analysis of distal colon revealed no discernible differences between untreated CR-Ψ and PBS mice, suggesting colonic recovery at 40 dpi (Fig. 1F, S1C). DSS treatment resulted in colonic inflammation as marked by loss of colonic crypt architecture, immune cell infiltration, sub-mucosal thickening and epithelial disruption, however, CR-Ψ mice displayed lower extent of damage in comparison to PBS mice (Fig. 1F, S1C). Together these results suggest that at 40 dpi colon shows recovery from the primary CR-mediated damage and inflammation, however, mice with a history of CR infection were protected from subsequent chemical induced colitis.

Since, PBS or CR-Ψ mice displayed similar weight loss during the 7 days of DSS treatment (Fig. 1B), and the differences in disease were observed on day 10 (7+3 model), we asked whether the differences observed were a consequence of lower disease during the DSS treatment or a better recovery after stopping DSS administration. To this end, we sacrificed PBS and CR-Ψ mice on day 7 of DSS treatment. Both PBS and CR-Ψ mice showed similar weight loss (Fig. S2A), however, CR-Ψ mice had longer colon (Fig. S2B-C) and lower colonic epithelial damage and inflammation as assessed in H&E-stained thin colon sections (Fig. S2D). This suggests that CR-Ψ mice are more resistant to DSS than the control mice.

### Co-housing did not affect CR-mediated DSS protection

Cage effect and gut microbiota have influence on the extent and severity of colitis caused by DSS^16,21^. To rule out cage effect and the influence that microbiota may have on the protection observed in CR-Ψ mice, we co-housed PBS and CR-Ψ mice post CR clearance for 20 days before the start of DSS treatment (Fig. 2A). As observed above, CR-Ψ mice continued to display lower susceptibility to colitis as evidenced by significant weight gain (Fig. 2B) and increased colon length post DSS treatment (Fig. 2C). Colonic inflammation assessed by H&E-stained distal colon sections revealed lower colonic crypt damage, immune cell infiltration and inflammation in co-housed CR-Ψ mice compared to co-housed PBS mice (Fig. 2D), suggesting that the observed protection observed in CR-Ψ mice is independent of cage effect and may not be a direct consequence of the long-term colonic microbiota changes associated with CR infection.

**Figure 2.**
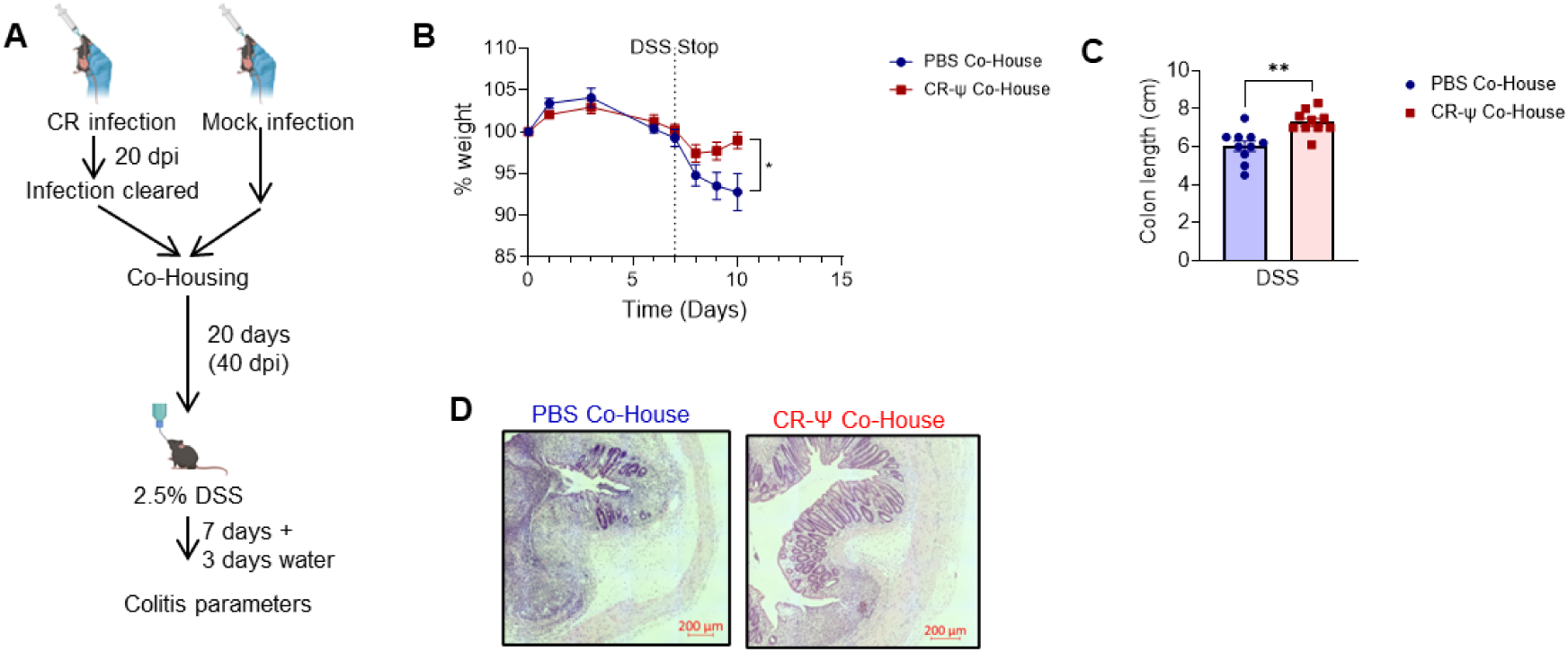
Protection from colitis is independent of cage effect. (A) Schematic representation of the Co-housing strategy. (B–D) Co-housed CR-Ψ mice retained protection from DSS-induced colitis, as indicated by improved weight (B), colon length (C) and histology (D). Scale 200 µm. Data shown are pooled values from 2 biological repeats with 5 mice in each group per experiment. P values were determined on data plotted as mean ± SEM using Student’s-t-test. ns: not significant; *p < 0.05; **p < 0.01.

### CR-induced disruption of tight junction is essential for protection from colitis

CR causes disruption of TJ, which is mediated by the T3SS effectors EspF and Map^4,22^. We hypothesised that disruption of the barrier function which triggers robust host immune response during the primary infection would be essential for training the epithelium and protection from the secondary DSS perturbation. To test this, we used a CR lacking Map and EspF (Δ*map*Δ*espF*). Mice with a history of Δ*map*Δ*espF* infection (Δ*map*Δ*espF*-Ψ) were equally susceptible to DSS-induced colitis as the control PBS mice. Δ*map*Δ*espF*-Ψ mice had similar weight to PBS mice post DSS treatment, which was significantly lower than CR-Ψ mice (Fig. 3A). Δ*map*Δ*espF*-Ψ mice displayed similar colonic shortening and colonic MPO levels to PBS mice post DSS treatment, suggesting similar extent of colonic damage and inflammation (Fig. 3B-D). Histological studies of distal colon revealed similar grade of loss of colonic crypt architecture, immune cell infiltration, sub-mucosal thickening, and epithelial disruption in PBS and Δ*map*Δ*espF*-Ψ mice (Fig. 3E). Overall, these results suggest that disruption of the TJ during CR infection is required for protection from subsequent colitis.

**Figure 3.**
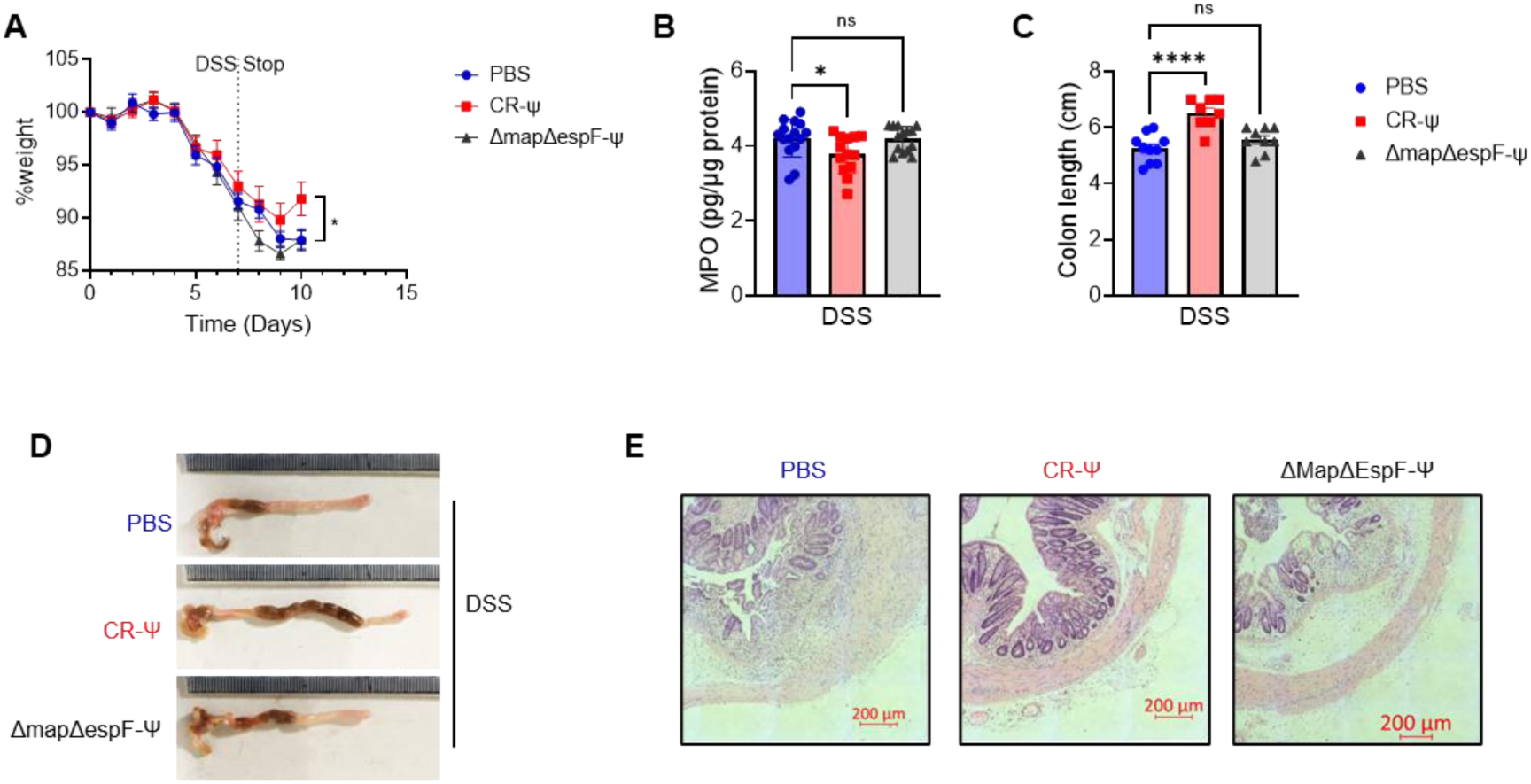
Protection from colitis requires optimal CR infection. (A) Weight change in CR-Ψ, Δ*map*Δ*espF*-Ψ, or PBS mice, post DSS treatment. (B) Colonic MPO. (C) Colon length. (D) Representative colon images. (E) Representative histology of distal colon revealed reduced crypt damage, immune cell infiltration, and epithelial disruption in CR-Ψ mice, while Δ*map*Δ*espF*-Ψ mice were comparable to PBS controls. Scale 200 µm. Data shown are pooled values from 2 biological repeats with 5 mice in each group per experiment. P values were determined on data plotted as mean ± SEM using Student’s-t-test in (A) and One-way ANOVA with Bonferroni post-test for multiple comparisons (B-C). ns: not significant, *p < 0.05; ****p < 0.0001.

### Mice with a history of CR infection had larger colonic T cell population

We investigated whether the protective imprint was associated with long-term changes in the colonic immune landscape—particularly T cell populations. We characterised the T cell profile in the colonic lamina propria at 40 dpi (Fig. 4A). Flow cytometry analysis using surface markers for T cells and its subtype revealed significantly more CD45+ cells, CD45+ CD3+ total T cell and CD4+ T cells in CR-Ψ mice compared to PBS or Δ*map*Δ*espF*-Ψ mice, suggesting long-term changes to T cell population post infection (Fig. 4B-D). Further characterisation of T cell subtypes revealed no significant differences in Treg and Th2 cell population (Fig. S3A-B), however, a significant increase in the number of Th1 and Th17 cells was observed in CR-Ψ mice compared to PBS or Δ*map*Δ*espF*-Ψ mice (Fig. 4E-F). Together, these results indicated that an expanded T cell population persist in the colonic lamina propria after CR clearance.

**Figure 4.**
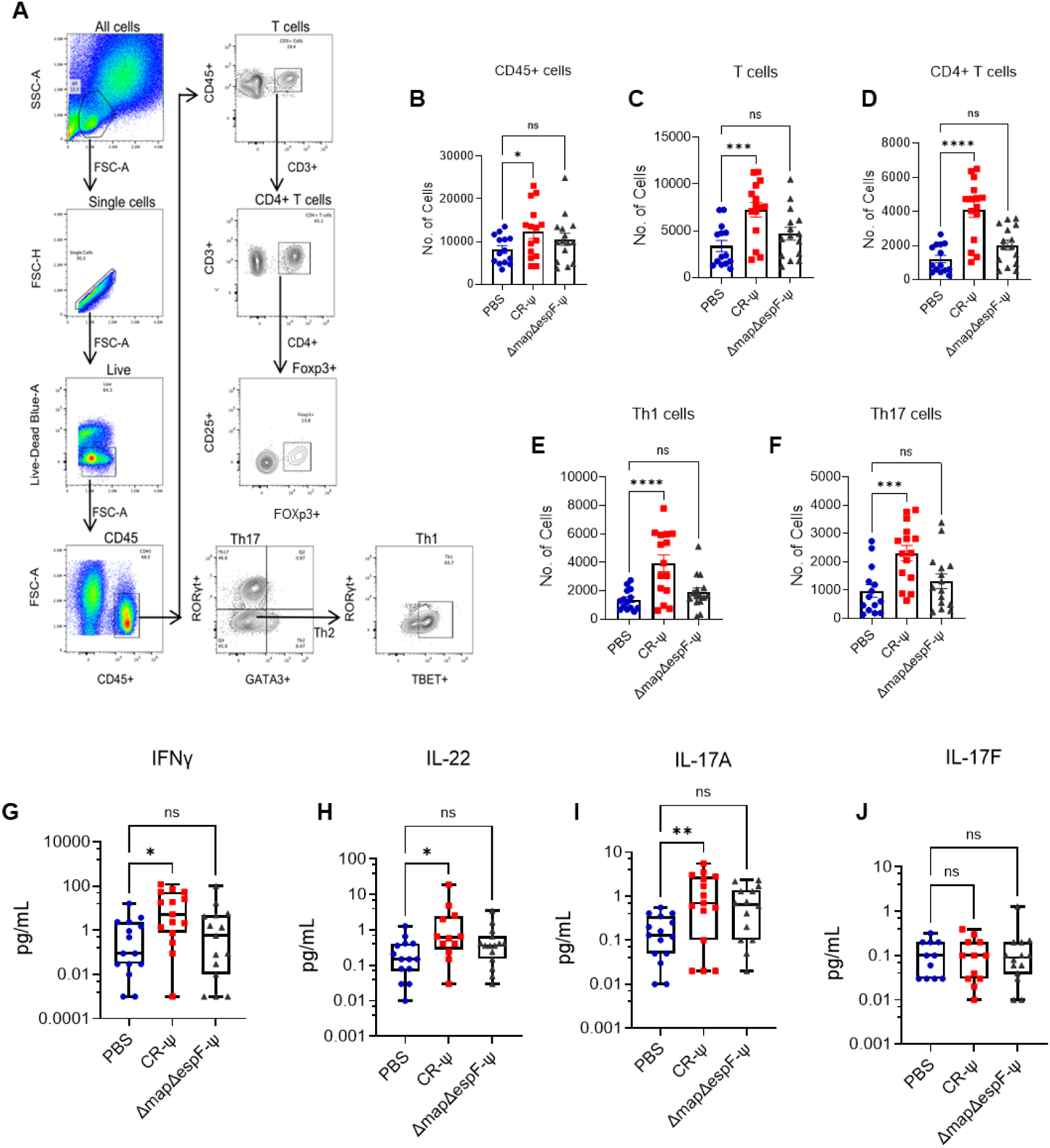
A history of wild-type CR infection results in long-term expansion of colonic T cells. (A) Gating strategy for flow cytometric analysis of lamina propria T cells. (B–F) Flow cytometric quantification of CD45⁺ leukocytes (B), CD45+ CD3+ total T cells (C), CD4⁺ T cells (D), Th1 cells (E) and Th17 cells (F) at 40 dpi. (G–J) Cytokine analysis of colonic explant culture supernatants at 40 dpi for IFNγ (G), IL-17A (H), IL-22 (I) and IL-17F (J). Data shown are pooled values from 3 biological repeats with 5 mice in each group per experiment. P values were determined on data plotted as mean ± SEM using One-way ANOVA with Bonferroni post-test for multiple comparisons. ns: not significant, *p < 0.05; **p < 0.01; ***p<0.001; ****p < 0.0001.

We next profiled the cytokines in the supernatant of colonic explant cultures at 40 dpi. CR-Ψ mice displayed significantly higher Th1 secreted IFNγ and IL-2 levels compared to PBS or Δ*map*Δ*espF*-Ψ mice (Fig. 4G, Fig. S4A). There were no significant differences in the type 2 cytokines IL-4, IL-5, IL-9 and IL-13 which is consistent with no observed differences in the abundance of Th2 cells between CR-Ψ and control mice (Fig. S4B-E). Moreover, no differences in the levels of Treg secreted IL-10 was observed (Fig. S4F). Similarly, all three groups of mice showed similar levels of TNF and IL-6 (Fig. S4G-H). However, significantly higher levels of Th17 mediated IL-22 and IL-17A was observed in CR-Ψ mice compared to PBS or Δ*map*Δ*espF*-Ψ mice, with no differences observed in the levels of IL-17F (Fig. 4H-J). Together, these results suggest that at 40 dpi there is an increased abundance of Th1 and Th17 cells the levels of their respective cytokines.

### Elevated levels of IL-17A results in protection from colitis

While mice lacking IFNγ are resistant to DSS-induced colitis and exogenous IFNγ treatment exacerbated disease severity^23,24^ and IL-22 is known to protect mice from DSS-induced colitis^12^, the role of IL-17A is inconclusive^14^. We therefore decided to investigate if IL-17A plays a role in protecting CR-Ψ mice from DSS. To this end we injected naïve mice with a total of three doses of recombinant IL-17A (rIL-17A) starting from 6 days before DSS treatment (Fig. 5A). Mice treated with rIL-17A displayed significant weight gain post DSS treatment (Fig. 5B) and longer colons (Fig. 5C-D). Moreover, histological analysis revealed lower inflammation and epithelial damage in mice treated with rIL-17A (Fig. 5E), phenocopying protection observed in CR-Ψ mice. These results indicated that elevated IL-17A may result in the protection observed in CR-Ψ mice.

**Figure 5.**
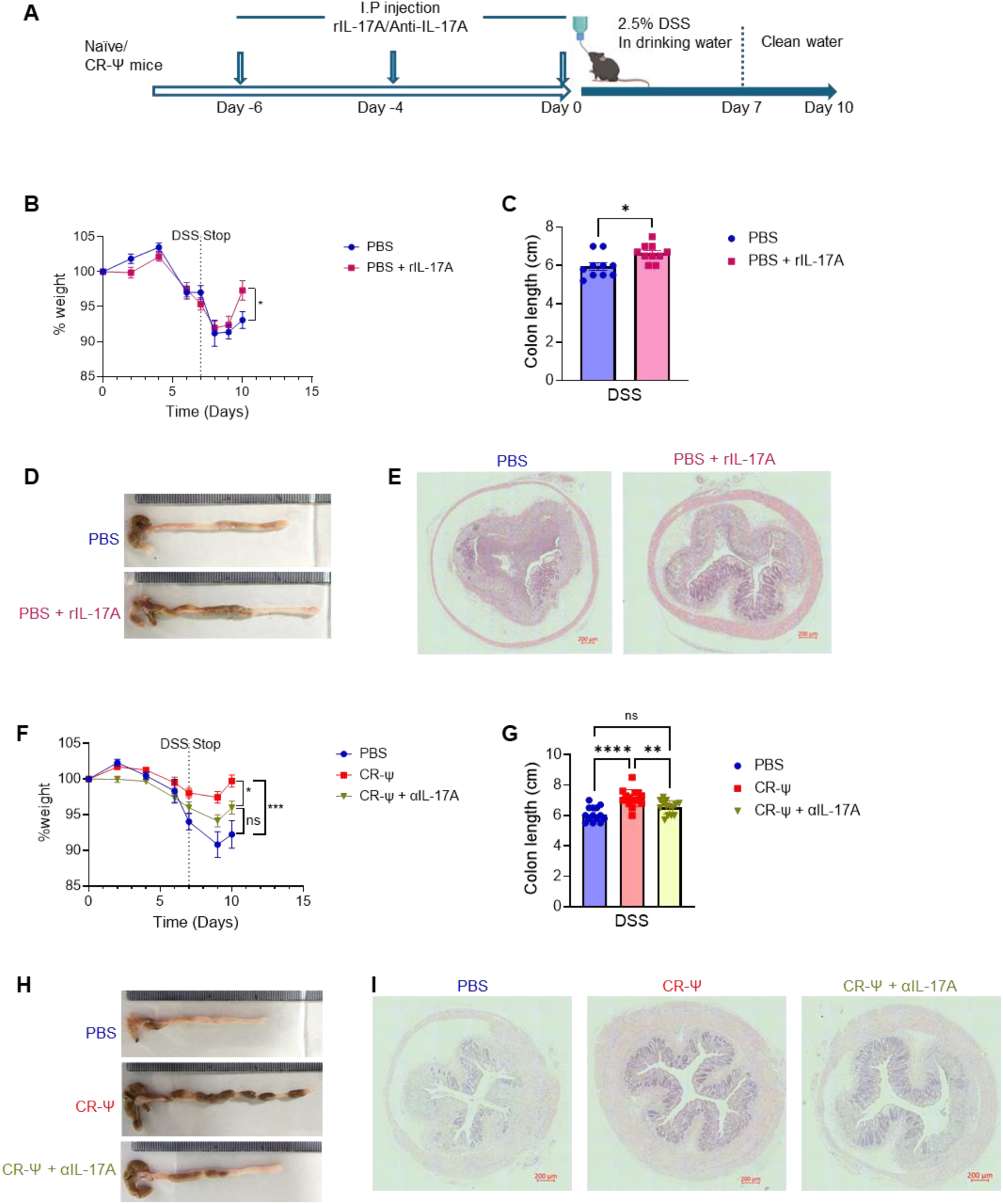
IL-17A is necessary and sufficient for protection from DSS-induced colitis. (A) Schematic representation of the treatment. (B–E) Naïve mice were pretreated with rIL-17A prior to DSS administration. rIL-17A-treated mice exhibited significantly improved weight (B), reduced colonic shortening (C-D) and lower colonic epithelial damage assessed by H&E-stained colonic sections (E). (F–I) CR-Ψ mice treated with anti-IL-17A neutralizing antibodies (αIL-17A) prior to DSS lost protection, as shown by lower weight (F) and reduced colon length (G-H), comparable to PBS controls. (I) Histological analysis revealed increased epithelial damage and inflammation in CR-Ψ + αIL-17A mice relative to untreated CR-Ψ controls. Scale 200 µm. Data shown are pooled values from 2 biological repeats for rIL-17A treatment group and 3 biological repeats for αIL-17A treatment group with 5 mice in each group per experiment. P values were determined on data plotted as mean ± SEM using Student’s-t-test in (B-C, F) and One-way ANOVA with Bonferroni post-test for multiple comparisons (G). ns: not significant; *p < 0.05; **p < 0.01; ****p < 0.0001.

To further establish the role of IL-17A in CR infection mediated protection from colitis, we treated CR-Ψ mice with anti-IL-17A antibodies prior to DSS treatment (Fig. 5A). CR-Ψ mice treated with anti-IL-17A lost significant weight compared to untreated CR-Ψ mice post DSS treatment, suggesting a loss of protection (Fig. 5F). CR-Ψ mice treated with anti-IL-17A displayed modest but non-significant difference in weight compared to PBS mice (Fig. 5F). However, no significant differences were observed in colon length between anti-IL-17-treated CR-Ψ and PBS mice, which was significantly shorter than CR-Ψ mice (Fig. 5G-H). Moreover, histological analysis revealed higher inflammation and epithelial damage in anti-IL-17-treated CR-Ψ compared to CR-Ψ mice (Fig. 5I). Together these results suggest that neutralisation of IL-17A prior to DSS treatment resulted in loss of protection observed in CR-Ψ mice, suggesting that in this context IL-17A confers colitis protection.

## Discussion

The interplay between microbial exposure and the immune system in shaping susceptibility to inflammatory diseases is complex and not well understood. Here, we demonstrate that a resolved intestinal infection with CR provides long-term protection against subsequent chemically induced colitis. These findings align with the “hygiene hypothesis,” which posits that reduced microbial exposures may contribute to rising immune-mediated diseases such as IBD^25^. Our study reveals a novel role for infection-induced immune imprinting in conferring protection against sterile colitis.

Infection-mediated protection against colitis required a robust inflammatory response during primary CR infection, induced following epithelial barrier disruption and mucosal inflammation. Mice infected with a Δ*map*Δ*espF* mutant which induces minimal crypt hyperplasia, reduced immune cell infiltration, and lower pro-inflammatory cytokine production^4^, failed to protect mice from subsequent colitis. Infection with Δ*map*Δ*espF* results in a marked reduction in IL-22 levels and its downstream epithelial targets^4^. Consistently, mice deficient in IL-22 succumb to wild-type CR infection but survive infection with Δ*espF* strains^26^. Interestingly, while at the peak of infection (8 dpi), recruitment of neutrophils, conventional dendritic cells, B cells, and CD4⁺ T cells are comparable between Δ*map*Δ*espF* and wild-type infections^4^, sustained expansion of CD4⁺ T cells after clearance appeared to require optimal TJ disruption and epithelial injury. Moreover, although primary infection with Δ*espF* strains can confer protection against homologous CR reinfection^27^, they fail to provide heterologous protection against chemically induced colitis. These findings suggest that the magnitude and nature of epithelial injury during initial infection are critical for establishing durable, protective mucosal immune reprogramming.

Although CR infection alters the gut microbiota^6^, our co-housing experiments indicate that immune imprinting, rather than long-term microbiota changes, underlies the observed protection. During infection, CR reduces microbial richness, with no changes in microbial diversity and alters the relative abundance of taxa such as Firmicutes and Enterobacteriaceae^5^. While such changes are associated with altered susceptibility to colitis^21^, the persistence of protection in co-housed mice with prior CR infection supports a dominant role for immune reprogramming over microbiota composition.

Mice displayed sustained increases in colonic Th1 and Th17 cells, along with elevated levels of their respective cytokines IL-2, IFNγ, and IL-22 and IL-17A post CR clearance. These T cells persisted in the lamina propria well beyond pathogen clearance and histological recovery, consistent with the concept of mucosal immune memory. CR infection is associated with long-term changes to the microenvironment of the colon resulting in elevated levels of IFNγ, epithelial MHCII expression and higher numbers of enteroendocrine cells^10^. CR infection generates long-lived tissue-resident IL-17A+ CD4+ memory T cells and trained ILC3s, which display enhanced IL-22 secretion and confer protection during reinfection^8,9^. Our findings expand this paradigm, showing that infection-primed T cells can also provide heterologous protection against sterile inflammatory insults.

Among the cytokines elevated in mice with a history of CR infection, IL-17A emerged as both necessary and sufficient for protection against chemically induced colitis. Administration of recombinant IL-17A significantly reduced disease severity following DSS treatment, while neutralisation of IL-17A in CR-experienced mice abrogated the protective effect. The role of IL-17A during DSS-induced colitis, however, has been historically controversial^14^. Early studies reported that IL-17A promotes acute mucosal inflammation, as *Il17a*-deficient mice exhibited protection from colitis^28^. Conversely, subsequent studies using *Il17a*-deficient mice or anti-IL-17A treatment revealed a protective function for IL-17A during DSS colitis, demonstrating that IL-17A supports epithelial repair by enhancing tight junction integrity, stimulating epithelial proliferation, and facilitating mucosal wound healing^15^. Moreover, in human studies, anti-IL-17A therapies that have shown efficacy in autoimmune conditions such as psoriasis paradoxically worsened disease outcomes in IBD patients, underscoring the complex and tissue-specific roles of IL-17A^14,29–31^. These contrasting effects likely reflect differences in the phase of disease (acute injury versus healing) and tissue context. During acute epithelial injury, IL-17A may exacerbate inflammation, whereas during the recovery phase, it appears critical for restoring barrier integrity and limiting further immune activation. Consistent with these observations, our findings suggest that IL-17A, when induced in a controlled and infection-resolved setting, fosters a mucosal environment more resilient to subsequent inflammatory insults.

Other cytokines such as IL-22 and IL-2 may act synergistically with IL-17A. The role of IL-22 is well established^12^, however, the role of IL-2 during DSS-induced colitis in mice is biphasic and dose-dependent. Low-dose IL-2 promotes immune tolerance and has shown benefit in DSS colitis models, while IL-2-deficient mice lacks adequate Treg populations and spontaneously develop systemic autoimmunity and colitis^32,33^. Although their individual roles remain complex, the combined presence of IL-22, IL-2, and IL-17A may contribute to a coordinated protective program in the post-infection mucosa.

Together, our results reveal a previously unrecognised benefit of resolved enteric infection in shaping long-term immune landscapes that protect against chemically induced colitis. These findings broaden our understanding of how prior microbial encounters influence future inflammatory responses and highlight the dual nature of IL-17A in health and disease. Future studies should explore how infection-primed T cells are maintained, whether similar protection can be achieved through vaccines or defined microbial exposures, and how these insights might inform novel strategies to prevent or treat IBD.

## Acknowledgements

VM thanks Imperial Early Career Researcher Institute ‘Seed for Success’ program for seed grant and Prof Sandhya S. Visweswariah for insightful discussions. We thank the staff of Central Biomedical Services, Imperial for assistance in maintenance of animals.

## References

1. Ananthakrishnan AN, Bernstein CN, Iliopoulos D, et al. Environmental triggers in ibd: A review of progress and evidence. Nature reviews Gastroenterology & hepatology 2018;15:39–49.

2. Collins JW, Keeney KM, Crepin VF, et al. Citrobacter rodentium: Infection, inflammation and the microbiota. Nat Rev Microbiol 2014;12:612–23.

3. Jordan S, Frankel G, Mishra V. Citrobacter rodentium. Trends Microbiol 2025.

4. Ruano-Gallego D, Sanchez-Garrido J, Kozik Z, et al. Type iii secretion system effectors form robust and flexible intracellular virulence networks. Science 2021;371.

5. He Y, Zhao J, Ma Y, et al. Citrobacter rodentium infection impairs dopamine metabolism and exacerbates the pathology of parkinson’s disease in mice. J Neuroinflammation 2024;21:153.

6. Hoffmann C, Hill DA, Minkah N, et al. Community-wide response of the gut microbiota to enteropathogenic citrobacter rodentium infection revealed by deep sequencing. Infect Immun 2009;77:4668–78.

7. Mullineaux-Sanders C, Sanchez-Garrido J, Hopkins EGD, et al. Citrobacter rodentium-host-microbiota interactions: Immunity, bioenergetics and metabolism. Nat Rev Microbiol 2019;17:701–15.

8. Serafini N, Jarade A, Surace L, et al. Trained ilc3 responses promote intestinal defense. Science 2022;375:859–63.

9. Bishu S, Hou G, El Zaatari M, et al. Citrobacter rodentium induces tissue-resident memory cd4(+) t cells. Infect Immun 2019;87.

10. Mullineaux-Sanders C, Kozik Z, Sanchez-Garrido J, et al. Citrobacter rodentium infection induces persistent molecular changes and interferon gamma-dependent major histocompatibility complex class ii expression in the colonic epithelium. mBio 2021;13:e0323321.

11. Park JH, Peyrin-Biroulet L, Eisenhut M, Shin JI. Ibd immunopathogenesis: A comprehensive review of inflammatory molecules. Autoimmunity reviews 2017;16:416–26.

12. Dudakov JA, Hanash AM, van den Brink MR. Interleukin-22: Immunobiology and pathology. Annu Rev Immunol 2015;33:747–85.

13. Sugimoto K, Ogawa A, Mizoguchi E, et al. Il-22 ameliorates intestinal inflammation in a mouse model of ulcerative colitis. The Journal of clinical investigation 2008;118:534–44.

14. Fauny M, Moulin D, D’Amico F, et al. Paradoxical gastrointestinal effects of interleukin-17 blockers. Ann Rheum Dis 2020;79:1132–8.

15. Zhou C, Wu D, Jawale C, et al. Divergent functions of il-17-family cytokines in dss colitis: Insights from a naturally-occurring human mutation in il-17f. Cytokine 2021;148:155715.

16. Chassaing B, Aitken JD, Malleshappa M, Vijay-Kumar M. Dextran sulfate sodium (dss)-induced colitis in mice. Curr Protoc Immunol 2014;104:15 25 1–15 25 14.

17. Yang C, Merlin D. Unveiling colitis: A journey through the dextran sodium sulfate-induced model. Inflammatory bowel diseases 2024;30:844–53.

18. Crepin VF, Collins JW, Habibzay M, Frankel G. Citrobacter rodentium mouse model of bacterial infection. Nat Protoc 2016;11:1851–76.

19. Kim E, Tran M, Sun Y, Huh JR. Isolation and analyses of lamina propria lymphocytes from mouse intestines. STAR protocols 2022;3:101366.

20. Hansberry DR, Shah K, Agarwal P, Agarwal N. Fecal myeloperoxidase as a biomarker for inflammatory bowel disease. Cureus 2017;9.

21. Ikeda E, Yamaguchi M, Kawabata S. Gut microbiota-mediated alleviation of dextran sulfate sodium-induced colitis in mice. Gastro Hep Adv 2024;3:461–70.

22. Nguyen M, Rizvi J, Hecht G. Expression of enteropathogenic escherichia coli map is significantly different than that of other type iii secreted effectors in vivo. Infection and Immunity 2015;83:130–7.

23. Langer V, Vivi E, Regensburger D, et al. Ifn-gamma drives inflammatory bowel disease pathogenesis through ve-cadherin-directed vascular barrier disruption. J Clin Invest 2019;129:4691–707.

24. Ito R, Shin-Ya M, Kishida T, et al. Interferon-gamma is causatively involved in experimental inflammatory bowel disease in mice. Clinical & Experimental Immunology 2006;146:330–8.

25. Segal AW. Making sense of the cause of crohn’s–a new look at an old disease. F1000Research 2016;5:2510.

26. Xia X, Liu Y, Hodgson A, et al. Espf is crucial for citrobacter rodentium-induced tight junction disruption and lethality in immunocompromised animals. PLoS Pathog 2019;15:e1007898.

27. Wang S, Xia X, Liu Y, Wan F. Oral administration with live attenuated citrobacter rodentium protects immunocompromised mice from lethal infection. Infect Immun 2022;90:e0019822.

28. Ito R, Kita M, Shin-Ya M, et al. Involvement of il-17a in the pathogenesis of dss-induced colitis in mice. Biochemical and biophysical research communications 2008;377:12–6.

29. Deng Z, Wang S, Wu C, Wang C. Il-17 inhibitor-associated inflammatory bowel disease: A study based on literature and database analysis. Frontiers in pharmacology 2023;14:1124628.

30. Hueber W, Sands BE, Lewitzky S, et al. Secukinumab, a human anti-il-17a monoclonal antibody, for moderate to severe crohn’s disease: Unexpected results of a randomised, double-blind placebo-controlled trial. Gut 2012;61:1693–700.

31. Grümme L, Dombret S, Knösel T, Skapenko A, Schulze-Koops H. Colitis induced by il-17a-inhibitors. Clinical Journal of Gastroenterology 2024;17:263–70.

32. Sund M, Xu LL, Rahman A, et al. Reduced susceptibility to dextran sulphate sodium-induced colitis in the interleukin-2 heterozygous (il-2+/–) mouse. Immunology 2005;114:554–64.

33. Tchitchek N, Tchoumba ON, Pires G, et al. Low-dose il-2 shapes a tolerogenic gut microbiota that improves autoimmunity and gut inflammation. JCI insight 2022;7:e159406.

