## Supplementary figures for "Enteric infection priming confers IL-17A–dependent protection from chemically-induced Colitis"

### Supplementary Figure 1

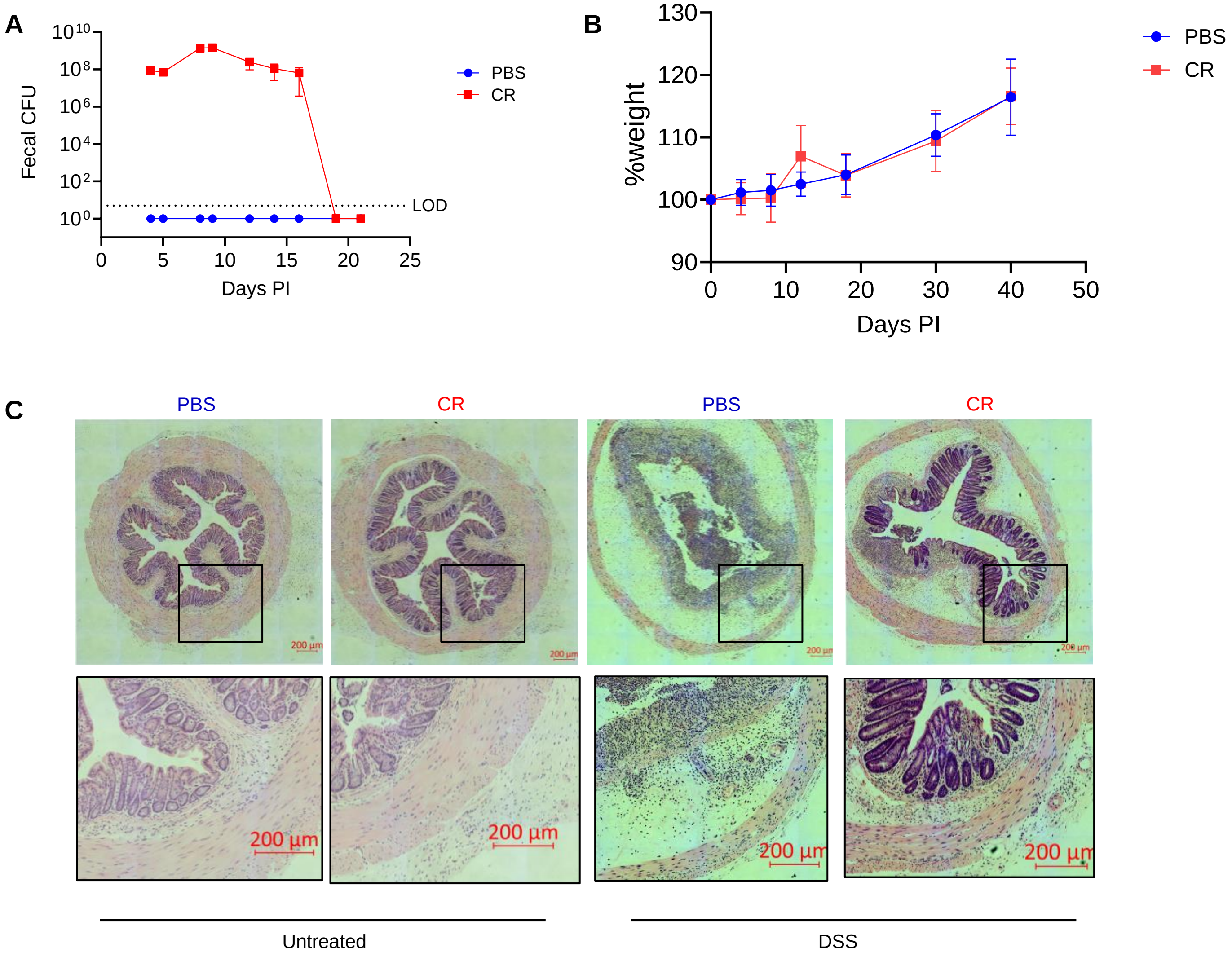


**Supplementary Figure 1. CR infection kinetics.** (A) Faecal CR shedding post oral infection, measured by CFU per gram of faeces. (B) Body weight of mice post-infection. (C) Representative H&E-stained colon sections. Scale 200 µm. Data shown are pooled values from 2 biological repeats with 5 mice in each group per experiment

**Supplementary Figure 2**

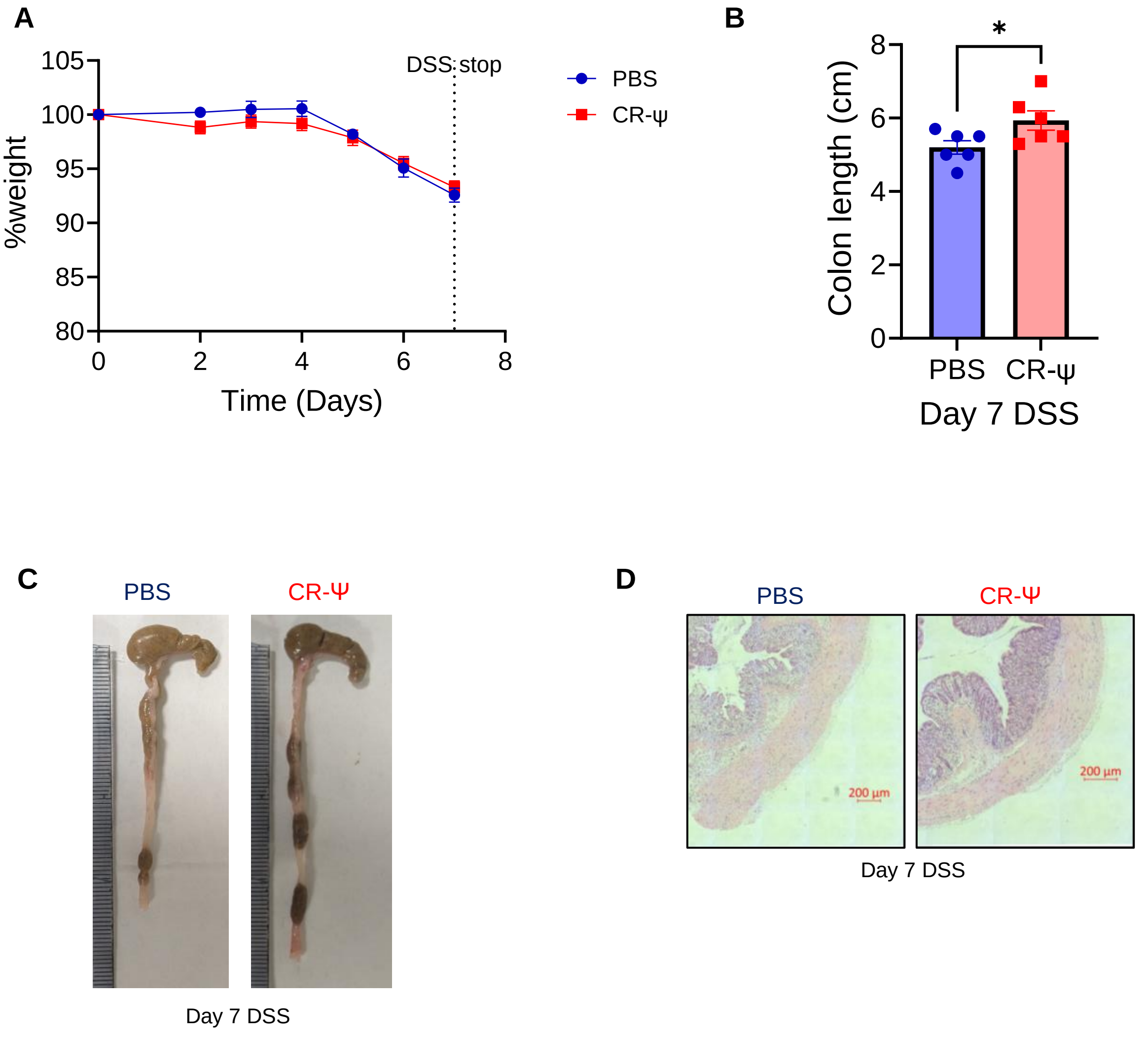


**Supplementary Figure 2. Disease parameters at day 7 of DSS treatment.** (A) Weight change during DSS treatment in PBS and CR-Ψ mice. (B-C) Colon length on day 7 was significantly longer in CR-Ψ mice, suggesting reduced early tissue damage. (D) Histological analysis at day 7 revealed reduced epithelial disruption and inflammatory infiltration in CR-Ψ mice compared to PBS controls. Data shown are pooled values from 2 biological repeats with 3 mice in each group per experiment. P values were determined on data plotted as mean ± SEM using Student's-t-test. *$p < 0.05$.

**Supplementary Figure 3**

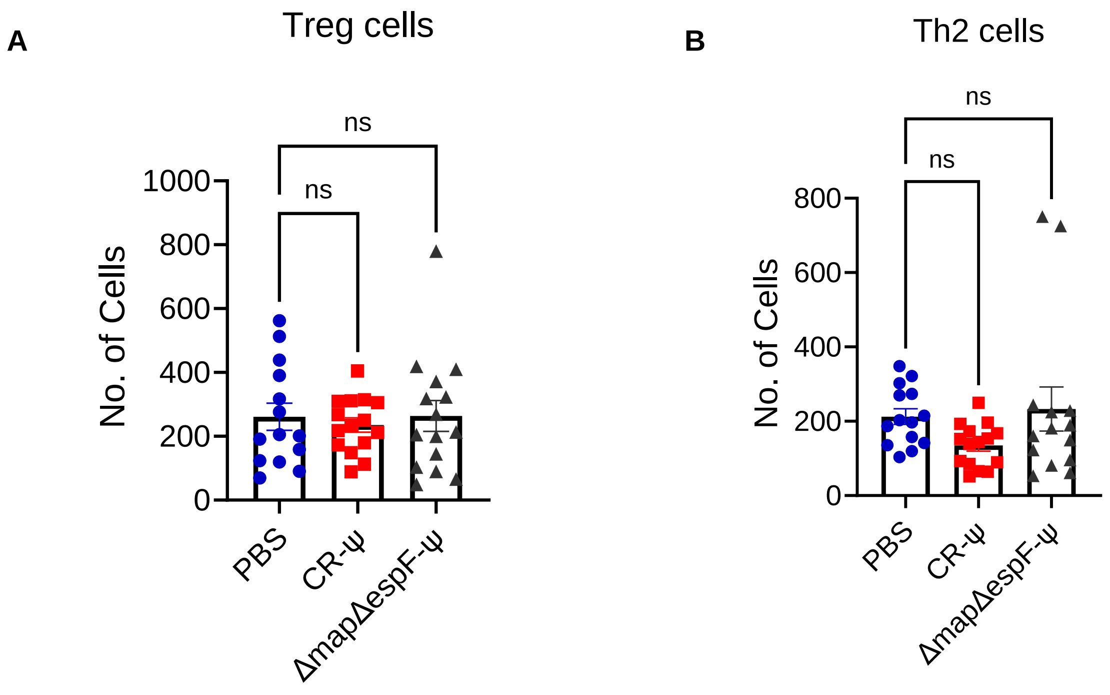


**Supplementary Figure 3. T cell profiling at 40 dpi.** (A-B) Number of Treg cells (A) and Th2 cells (B) in lamina propria. Data shown are pooled values from 3 biological repeats with 5 mice in each group per experiment. P values were determined on data plotted as mean ± SEM using One-way ANOVA with Bonferroni post-test for multiple comparisons. ns: not significant.

### Supplementary Figure 4

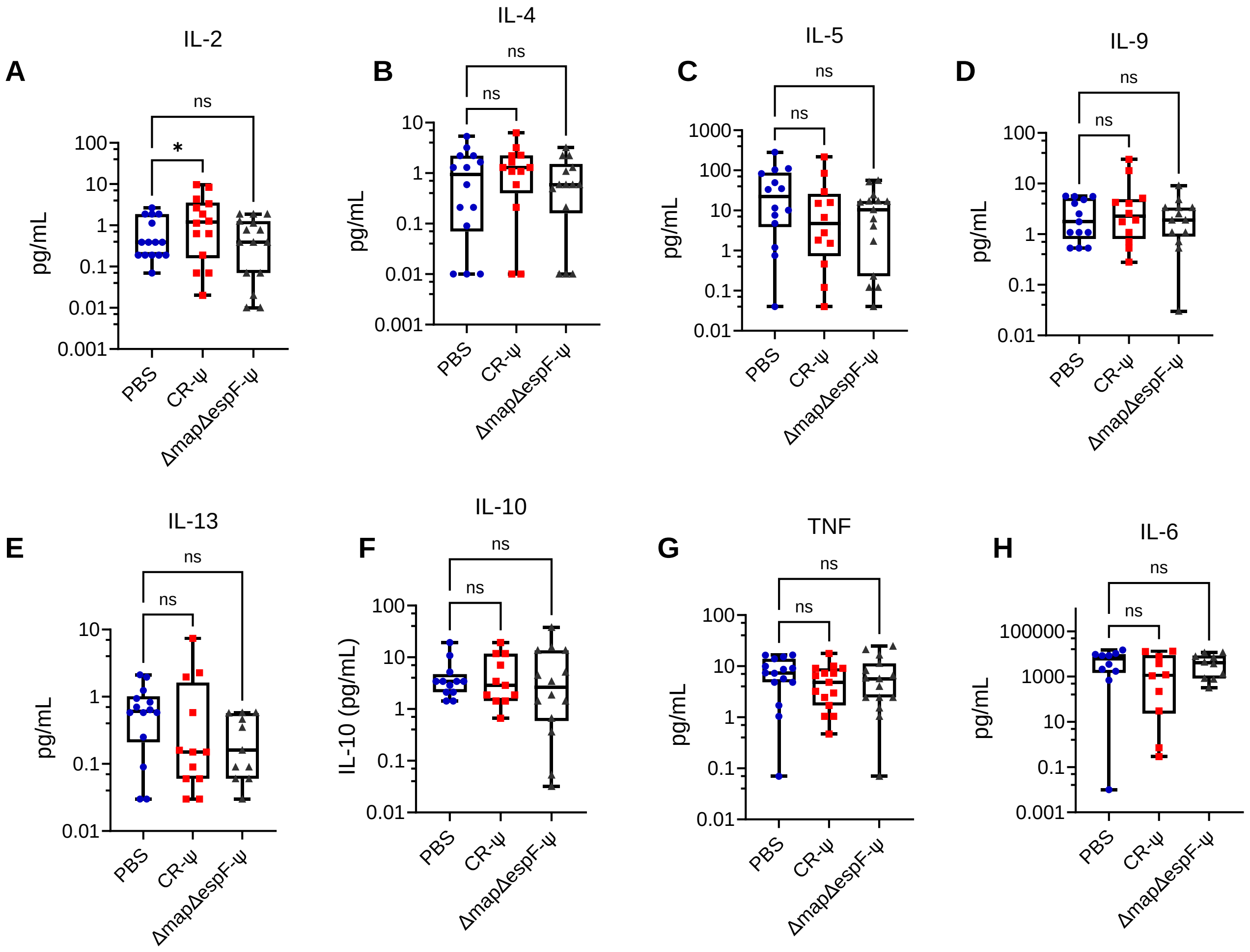


**Supplementary Figure 4. Cytokine profiling at 40 dpi.** (A–I) Cytokine profiling of colon explants showed for IL-2 (A), IL-4 (B), IL-5 (C), IL-9 (D), IL-13 (E), IL-10 (F), TNF (G), and IL-6 (H). Data shown are pooled values from 3 biological repeats with 5 mice in each group per experiment. P values were determined on data plotted as mean ± SEM using One-way ANOVA with Bonferroni post-test for multiple comparisons. ns: not significant; *p < 0.05.
